# Relative matching using low coverage sequencing

**DOI:** 10.1101/2020.09.09.289322

**Authors:** Ella Petter, Regev Schweiger, Bar Shahino, Tal Shor, Malka Aker, Lior Almog, Daphna Weissglas-Volkov, Yoav Naveh, Oron Navon, Shai Carmi, Jeremiah H. Li, Tomaz Berisa, Joseph K. Pickrell, Yaniv Erlich

**Affiliations:** MyHeritage LTD, Or Yehuda, Israel; Department of Public Health, School of Medicine, Hebrew University, Jerusalem, Israel; Gencove Inc, New York, NY

## Abstract

Finding familial relatives using DNA has multiple applications, in genetic genealogy, population genetics, and forensics. So far, most relative matching algorithms rely on detecting identity-by-descent (IBD) segments with high quality genotype data. Recently, low coverage sequencing (LCS) has received growing attention as a promising cost-effective method to ascertain genomic information. However, with higher error rates, it is unclear whether existing IBD detection can work on LCS datasets. Here, we developed and tested a framework for relative matching using sequencing with 1× coverage (1×LCS). We started by exploring the error characteristics of this method compared to array data. Our results show that after some optimization 1×LCS can exhibit the same genotyping discordance rates as the discordance between two array platforms. Using this observation, we developed a hybrid framework for relative matching and tuned this framework with >2,700 pairs of confirmed genealogical relatives that were genotyped using heterogenous datasets. We then obtained array and 1×LCS on 19 samples and use our framework to find relatives in a database of over 3 million individuals. The total length of shared segments obtained by 1×LCS was virtually indistinguishable to genotyping arrays for matches with a total sharing >200cM (second cousins or closer). For more distant relatives, as long as those were detected by both technologies, the total length obtained by LCS and by genotyping arrays was highly correlated, with no evidence of over- or underestimation. Taken together, our results show that 1×LCS can be a valid alternative to arrays for relative matching, opening the possibility for further democratization of genomic data.

## Introduction

Detection of familial relationship from DNA data is one of the most popular features in consumer genomics, which has gained tremendous popularity over the past few years. As of 2020, over 30 million people have taken a consumer genomics test, which enabled the creation of large scale genetic databases (Regalado, 2019). While some genetic direct to consumer products focus on health information, it appears most of the participants have taken tests that focus on genealogy products (Yuan et al. 2018; Khan & Mittelman 2018). These usually include a DNA relative matching feature that provides the costumers with a report revealing all of their genetic matches in the database in a sorted order, which typically go beyond second and third cousins. As most individuals do not maintain familial relationships with such distant relatives, finding genealogical relatives from genetic data has become a central tool in genealogical quests. Recently, this feature gained tremendous popularity in investigative genetic genealogy, where it has been used to solve hundreds of violent crimes and identify unclaimed bodies by finding distant relatives of crime-scene samples (Ney et al. 2020; Erlich et al. 2018). Finally, finding genealogical relatives from DNA also have applications in medical genetics (Speed & Balding 2015) and population genetics (Atzmon et al. 2010).

Finding genealogical relatives from DNA typically relies on algorithms to detect identity-by-descent (IBD) segments, identical segments of DNA inherited from a common ancestor (Browning & Browning 2012; Ramstetter et al. 2017). The IBD segments appear as a series of consecutive genotypes, where at least one allele of each marker is shared between a pair of individuals. Thus, many IBD detection techniques work on phased haplotypes, where the genotypes also have been imputed to avoid missing markers. Then, many algorithms use a seed and extend approach, in which small identical regions in a pair of haplotypes are quickly identified and are then extended until the IBD segment is over and the haplotypes are no longer identical.

IBD detection algorithm in different forms have mainly been applied on dense genotyping arrays or whole-genome sequence data (Huff et al. 2011; Li et al. 2014; Ramstetter et al. 2018; Zeng et al. 2016). These data sources exhibit very low genotyping errors rates of less than 0.1%, which can be easily overcome by most IBD detection algorithms by extending the IBD segments while tolerating a small number of errors. Empirical studies have reported that it is possible to find distant relatives from these datasets with quite high reliability. For example, 80% of the third cousins can be identified using these IBD segment matches and about 50% of all fourth cousins (Huff et al. 2011). These numbers are close to the theoretical limit, which is dictated by the probability of two individuals with a particular familial relationship sharing an IBD segment.

Here, we focus on the possibility of utilizing 1× low coverage sequencing (1×LCS) to find genetic relatives via IBD detection. LCS harnesses the power of high throughput sequencing with its great scaling trajectory while keeping the sequencing costs at the range that is affordable for consumer genomics. Due to that, LCS has received growing attention in the past few years, with applications in genome-wide association studies, ethnicity estimates, and predictions of polygenic risk scores (Li et al. 2020; Wasik et al. 2019; Pasaniuc et al. 2012). However, 1× sequencing coverage necessarily entails that about half of the genotypes reflect only one out of the two possible alleles in the autosomal regions. While imputation can enhance these calls, previous research reported that the discordance of 1×LCS after imputation is around 5% for common markers and samples of European ethnicity (Li et al. 2020), about one order of magnitude worse than the accuracy of SNPs reported by genotyping arrays (LaFramboise 2009), posing a challenge for IBD algorithms.

To address that we developed a hybrid tolerance method that can tolerate errors in IBD segments while maintaining some strictness to keep adequate levels of specificity. We then inspected the performance of our method with 1×LCS and compared it to IBD matching results of common arrays. Finally, we conclude that 1×LCS data can produce IBD matching result of similar quality to arrays.

## Results

### Collecting 1×LCS and baseline array data

We collected DNA samples from 19 participants that agreed to participate in this study (**Table 1**). This set of participants were mainly of European and Central Asian ethnicities and they contributed DNA mainly using the standard MyHeritage kit which relies on a buccal swab tip applicator that is immersed in a DNA extraction buffer and shipped at room temperature to the laboratories (see **Methods**).

**Table 1.** description of the 19 samples used in the experiment. Each individual was both genotyped by a chip array and sequenced and down-sampled to 1× coverage.

| Sample Number | Sex | Main ethnicity groups | Number of mapped reads | Downsampled autosomal coverage | Array Type |
| --- | --- | --- | --- | --- | --- |
| 1 | Female | Scandinavian | 19,515,311 | 0.941 | MH GSA |
| 2 | Male | Northwest European | 19,577,395 | 0.957 | MH GSA |
| 3 | Female | Scandinavian | 19,759,562 | 0.952 | MH GSA |
| 4 | Female | Greek, Italian | 19,738,121 | 0.951 | MH GSA |
| 5 | Female | British | 19,834,583 | 0.955 | MH GSA |
| 6 | Female | Scandinavian, North Western European | 19,894,535 | 0.960 | MH GSA |
| 7 | Male | Ashkenazi Jewish | 19,626,660 | 0.957 | 23andMe Version3 OmniExpress |
| 8 | Male | Ashkenazi Jewish and Central Asian | 19,451,300 | 0.949 | MH OmniExpress |
| 9 | Male | Irish, Scottish, and Welsh | 18,929,183 | 0.918 | MH OmniExpress |
| 10 | Female | Ashkenazi Jewish | 19,150,803 | 0.917 | MH OmniExpress |
| 11 | Female | Finnish, West Russian | 16,683,352 | 0.798 | MH OmniExpress |
| 12 | Female | British and Irish | 16,061,301 | 0.766 | MH OmniExpress |
| 13 | Male | Scandinavian | 19,650,579 | 0.962 | MH OmniExpress |
| 14 | Male | British and Scandinavian | 19,818,210 | 0.971 | MH OmniExpress |
| 15 | Male | Finnish, West russian | 19,685,215 | 0.965 | MH OmniExpress |
| 16 | Male | North Western European | 19,843,960 | 0.974 | MH OmniExpress |
| 17 | Female | British | 19,765,686 | 0.953 | MH OmniExpress |
| 18 | Female | Scandinavian | 19,710,889 | 0.945 | MH OmniExpress |
| 19 | Female | Scandinavian | 19,862,266 | 0.956 | MH OmniExpress |

We subjected the samples to DNA sequencing of paired-end 150bp reads using the BGI CompleteGenomics T10 DNBseq technology. Previous studies showed that the accuracy of this technology is similar to Illumina sequencing (Kim et al. 2020; Senabouth et al. 2020). We computationally downsampled the sequence reads to obtain 1×LCS by randomly selecting 10^7^ paired-end sequence reads from the FASTQ file of each sample. Next, we aligned the reads to the human genome and produced a VCF file using the automatic pipeline of Gencove (**Methods**). On average, 96.5% (19.29M±0.31M) of the reads aligned to the human genome, creating an average coverage of 0.934× (SD=0.055) of the autosome (**Table 1**). Each imputed VCF file had on average 21.2M genotypes that were based on observing at least a single read and 17.8M genotypes that were based on a fully imputed value without a single sequence read. Similar percentage were observed for the entire genome (**Supplementary Figure 1**).

To assess the accuracy of the LCS results, we ran a genome-wide array on each sample, in addition to low coverage sequencing. The genotyping arrays were then processed in the standard MyHeritage pipeline that has been used to process over 3 million samples so far. We processed 12 out of the 19 samples with the MyHeritage-customized OmniExpress array technology that genotypes 702,442 variants across the genome and was mainly developed for genetic genealogy purposes; and one with a 23andMe v3 OmniExpress array with 960,614 variants. The other 6 samples were processed with the MyHeritage-customized GSA array. This array genotypes 671,920 variants across the genome and includes tens of thousands low frequency variants that are associated with various health conditions.

### The genotyping accuracy of 1×LCS compared to arrays

We found that the imputation panel plays an important role in the accuracy of the 1×LCS genotyping results. At first, we imputed the samples using the 1000Genomes panel of Gencove, which is the default scheme that was used in previous studies. One advantage of this panel is that it contains very rare variants and therefore we hypothesized that it can provide better concordance with the GSA array that is enriched with these variants. We then calculated the genotype concordance between the ~700,000 genotypes inspected by the array and the genotypes of 1×LCS technology of the same individual that were imputed using the 1000Genomes. To this end, we measured the non-reference concordance (NRC) across genotypes. This measure assesses the concordance of all sites where at least one allele in either technology displayed the minor allele (see **Methods**). On average, we found an NRC of 94.66% across the samples (**Supplementary figure 2**). Next, we tried to impute using the Haplotype Reference Constrictum (HRC) panel that has approximately 10x more genome but not ultra-rare alleles (S et al. 2016). Importantly, the average NRC increased to 95.37%, reducing the errors in non-reference genotypes by 10% (**Figure 1A**, Fisher’s exact test: p<1e-100) with this imputation panel. Due to the improved results, in all of our next experiments, we switched to the HRC imputation to assess 1×LCS and the arrays.

**Figure 1.**
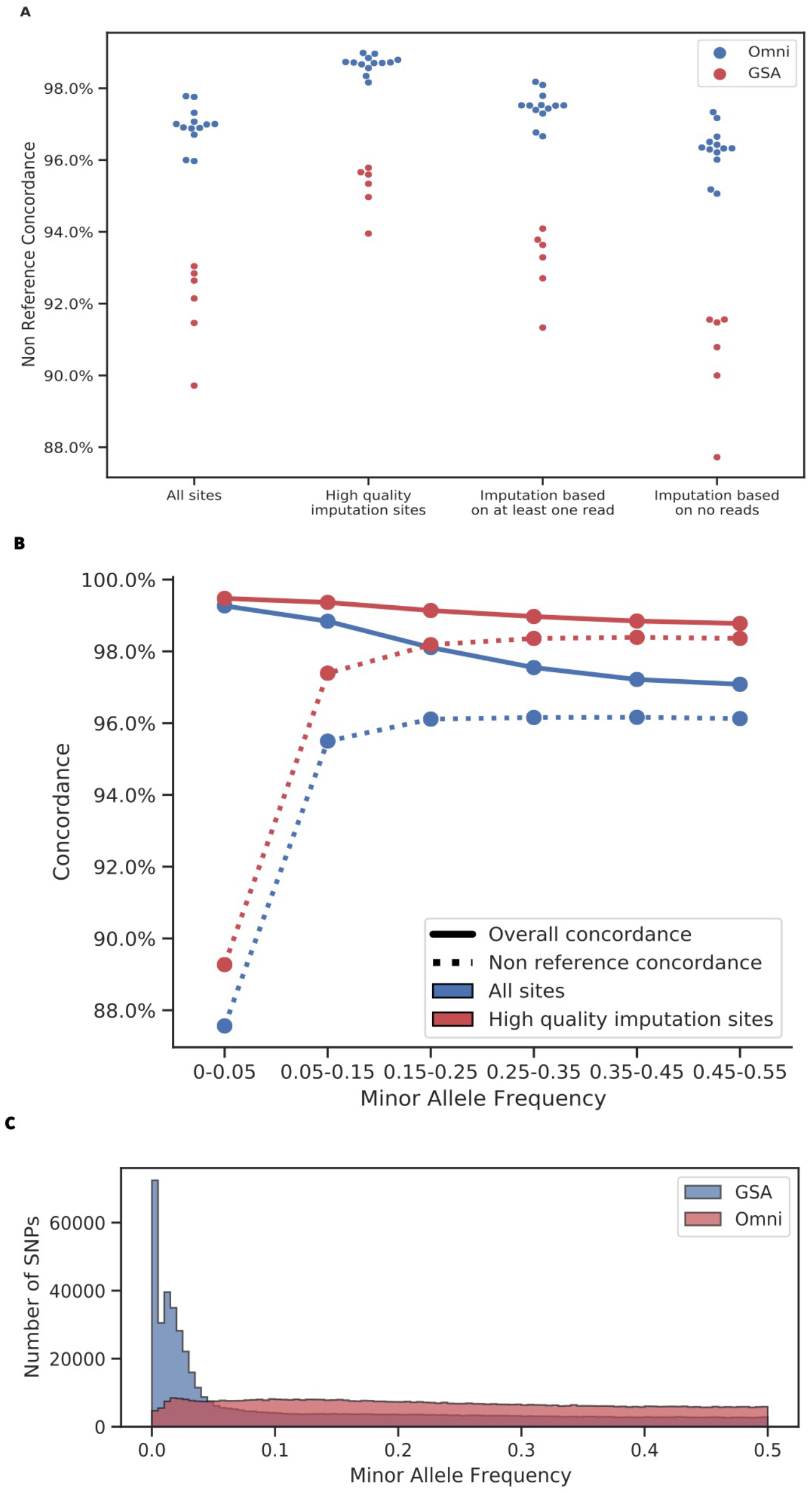
(A) NRC obtained for the 19 l×LCS samples compared to their matching array, under imputation based on the HRC reference panel. (B) Average concordance and non-reference concordance between l×LCS and arrays. In blue, considering all VCF sites, in red, only high quality imputation sites. (C) Histograms of minor allele frequencies of SNPs on the MyHeritage Omni and GSA arrays.

We also noted that the presence of even a single sequence read can significantly improve the accuracy of the genotypes. When the imputation was supported by at least one sequence read, the rate of errors in non-reference sites (1-NRC) was 3.9%. When the genotype was fully imputed without a single sequence read, the error rate in non-reference sites was 5.6%, 1.43x more compared to sites that were supported by a single read (Fisher’s exact test: p<1e-100). This observation suggests that LCS with around 1x coverage creates a trade-off between cost and quality, since the majority of SNPs are covered by at least a single sequence read, which improves the quality of the data, while still minimizing the per sample sequencing costs. A complementary analysis was also performed on SNPs where the imputation pipeline reported genotype quality higher than 0.9 (see **Methods)**. We refer to these SNPs as “high quality imputation SNPs”. These SNPs had an average NRC of 97.5%.

We found a notable difference between the concordance of 1×LCS compared to Omni and GSA arrays. The NRC showed a bimodal distribution and was better by 4% with the OmniExpress array compared to GSA in all conditions. To further probe into the lower concordance with the GSA arrays, we performed a concordance analysis stratified by the minor allele frequencies (**Figure 1B**). This time, we calculated both the NRC concordance and also the total concordance, which is not conditioned on the presence of a minor allele. The average concordance remained high for all allele frequencies, but the NRC was substantially lower for rare sites, as expected. This behaviour replicates results that were reported by a previous study (Li et al. 2020) and can be explained by a difficulty in imputing rare sites correctly. The MyHeritage GSA array has substantially more rare genetic variants than the Omni array (**Figure 1C**). Thus, we concluded that the lower NRC of 1×LCS for the GSA arrays is due to the enrichment of these SNPs with lower MAFs in GSA arrays.

We sought to establish the imputation concordance of the two array platforms to contextualize the concordance between 1×LCS to arrays. We considered two scenarios: (a) inter-array concordance, in which the same sample is genotyped by two array platforms, namely MyHeritage GSA and MyHeritage Omni arrays (b) intra-array concordance, in which the same sample is genotyped by the same array platform in two independent times. In both scenarios, we analyzed 1.5M common markers that were readily accessible for fast compute. For the inter-array comparisons, we obtained 1500 samples from the MyHeritage database of users who were tested with both a MyHeritage Omni and a MyHeritage GSA arrays. We imputed these samples using the HRC panel and analyzed their genotyping concordance. This process yielded an inter-array NRC of 96.27% on average. For the intra-array concordance, we ascertained 1195 samples of the MyHeritage database who were genotyped twice by Illumina Omni arrays and subjected them to HRC imputation as well. This process yielded an inter-array NRC concordance of 99.72% on average.

The inter- and intra-array comparisons highlight two phenomena: first, that the discordance between 1×LCS and arrays is within the discordance rates of switching between two array platforms, such as Omni and GSA. Second, the high discordance rate for inter-array comparisons stems from differences in array platforms and not the repeatability of the same array genotyping, which is quite high. Taken together, these results suggest that relative matching using 1×LCS is a private instance of a general problem of relative matching using heterogenous genomic datasets, such as two array platforms.

### Optimizing the matching algorithm using heterogeneous genomic data with confirmed genealogical relationships

Motivated by our observations, we moved to create an IBD matching algorithm for heterogenous datasets. The MyHeritage IBD detection pipeline uses a massively parallel implementation of a GERMLINE-like seed-and-extend algorithm (Gusev et al. 2009) that can tolerate errors in the extension part. A previous analysis has suggested that the best performance of such seed-and-extend algorithms is achieved with a tolerance of up to 2 errors in homozygous sites and 2 errors in heterozygous sites (Ramstetter et al. 2018; Huff et al. 2011). However, this error tolerance was determined using a homogenous dataset, which is not applicable for our situation.

To tune the error tolerance, we sought for a large collection of heterogenous datasets with confirmed genealogical relationships. However, generating a large number of 1×LCS datasets for such relatives is both logistically challenging and cost prohibitive. As an alternative, we focused on a set 2,711 pairs of relatives with validated genealogical relationships from parent-child to fourth cousins. Importantly, one person in each pair was tested with an Omni array whereas the other person was tested with a GSA array. Our working hypothesis was that since GSA and Omni have similar patterns of discordance as GSA/Omni vs 1×LCS, optimizing the error tolerance of our pipeline with these examples will also improve the performance of the 1×LCS relative matching.

As expected, we experienced poor performance of the IBD pipeline for these 2,711 pairs of relatives using the naïve error tolerance that was tuned for homogenous datasets. For example, using the naïve error tolerance, we detected 37 segments of IBD for parent-child pairs instead of exactly 22 segments as required by Mendalian inheritance. This indicates that the pipeline breaks IBD segments due to a large number of errors. In addition, these pairs shared only 3230cM on average, instead of >3500cM in previous studies with homogenous datasets, again meaning that some shared segments are missed.

To work with heterogenous datasets, we developed a hybrid approach for detecting IBD regions and performed a grid search to find the optimal error tolerance using true relatives (**Table 2**). Our hybrid approach retains the set of IBD segments in a pair only if at least one of the segments contains more than 6cM without a single error. We hypothesized that the evidence of at least one “high confidence” IBD segment increases the chance that the other segments are true as well and not just the product of many errors. Using this strategy, we performed a grid search for various heterozygous and homozygous parameters and evaluated its performance of detecting true relatives in the heterogeneous dataset. When using this framework, we noticed that only with 4 heterozygous SNPs and up to 4 homozygous SNPs the total IBD and the number of segments of Omni-GSA comparisons recapitulated the expected results from homogenous array datasets without inflating too much the IBD in homogenous datasets.

**Table 2.**
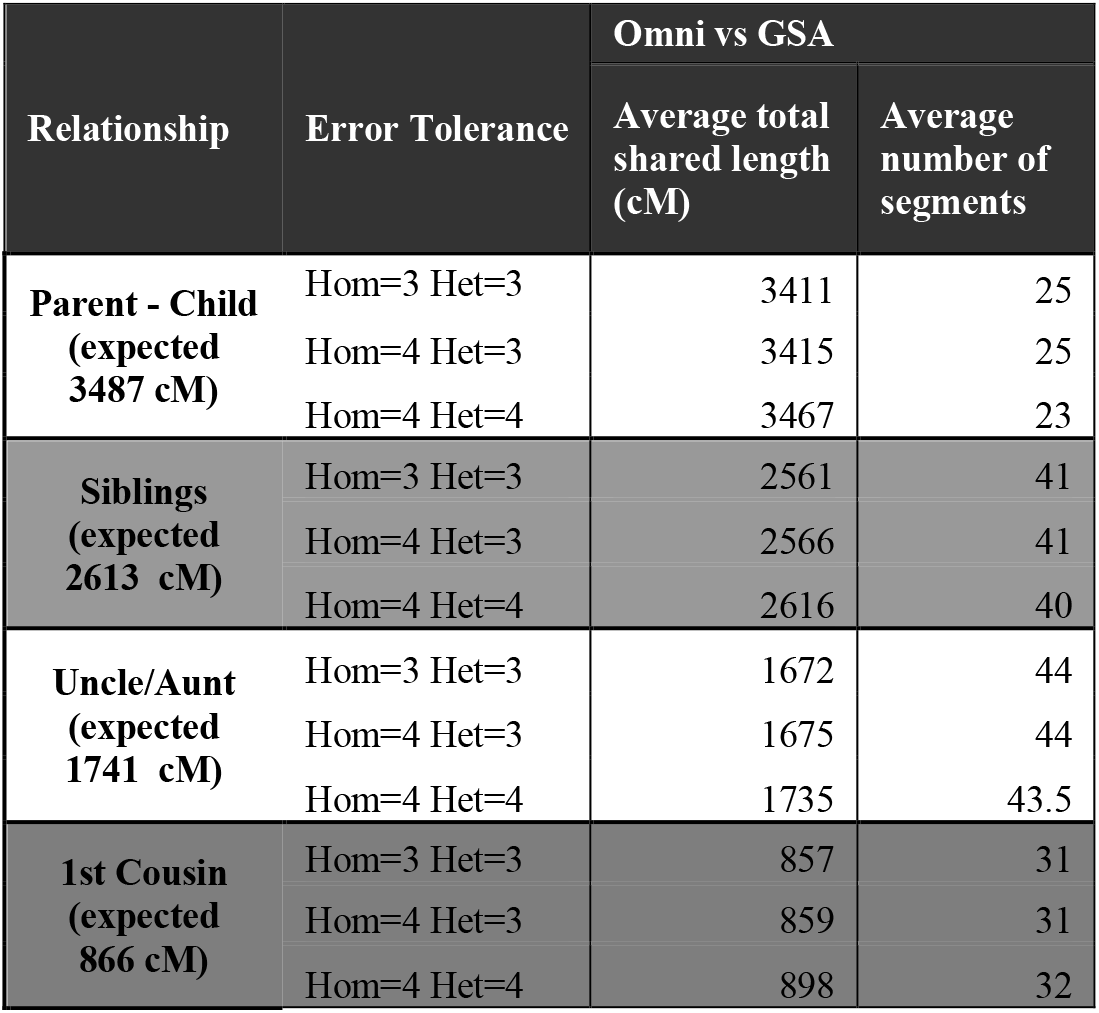
Average total shared length (cM) and number of segments found for matches in different relationships and array technologies, under different levels of error tolerance. Expected values are taken from The Shared cM Project (Bettinger & Perl 2018).

We were concerned that our permissive error tolerance can inflate the rate of false IBD segments. To measure this rate, we attained trios from our data and inspected the set of genetic relatives for each trio. Generally, we expect that if the child of the trio shares IBD segments with a certain individual outside of the trio, then one of the parents will also share IBD with this individual. Thus, any case where a child shares IBD with an individual in the database, while neither of their parent does, can indicate a false-positive event for the child or a false negative event for the parent. Therefore, this rate is a conservative measure on the false positive rate. We tested the child-only IBD sharing with our hybrid strategy using the naïve error tolerance (hom=2, het=2) and the new error permissive parameters (hom=4, het=4). We saw only a modest jump in the child-only IBD sharing, from 20.8% in the naïve error tolerance to 22.1% with the permissive rate. This means that the permissive error parameter with the hybrid strategy of a 6cM no-error filter adds only a small inflation in the false positive rate. This result led us to choose the permissive parameters of 4 homozygous and 4 heterozygous error, combined with the filtering step for 1×LCS.

### The total IBD between relatives by 1×LCS successfully replicates array results

Encouraged by our results with the GSA versus Omni, we applied the permissive relative matching algorithm on both the 1×LCS and the array based datasets and compared our 19 samples to 3.5 million individuals in our database for each technology.

As a first line of analysis, we measured whether there are systematic differences in the total IBD sharing between 1×LCS and arrays, focusing on close relatives above 200cM (second cousins and closer). These relatives are the most important for genetic genealogy and users are highly sensitive for errors in these results. For each pair of close relatives, we summed the total length of the IBD segments and compared the results between the array and 1×LCS. We then pooled the relatives from all 19 samples, where at least one of the technologies reported a shared length of at least 200cM and performed a linear regression of the total IBD using the 1×LCS as the dependent variable and the array results as an independent variable with zero intercept.

Our results indicate that the total IBD sharing for close relatives is highly similar between 1×LCS and arrays. Inspecting 67 close relatives yielded a regression slope of 0.998 (95% bootstrap CI: 0.995-1.001, see **Methods**), tightly following the diagonal (**Figure 2A**). We also found a correlation of R=0.99 for these relatives, further indicating the similarity.

Next, we repeated the same analysis for distant relatives (<200cM). By separating the close and distant relatives for the linear regression, we mitigated the effect of outliers from close relatives on the procedure. As we were interested in detecting systematic differences in total IBD sharing, we excluded any relatives that were detected by one but not the other technology (these will be discussed in the next section). Again, we found that the even for distant relatives, the total IBD sharing is quite similar between 1×LCS and arrays. When considering 141,127 distant relatives (<200cM) across all 19 samples, the slope of the regression line was 1.052 (95% bootstrap CI: 1.022-1.025, see **Methods**), very close to the diagonal line (**Figure 2B**). When we repeated this experiment for each of the 19 samples independently, the average slope of the regression line was 1.002, maintaining similar result for the samples separately. This indicates that 1×LCS does not tend to over- or underestimate the total IBD sharing compared to arrays. In addition, the correlation of the total IBD sharing between technologies for the distant relatives was R=0.87. Again, these results indicate that in general, the 1×LCS provides similar results to arrays.

**Figure 2.**
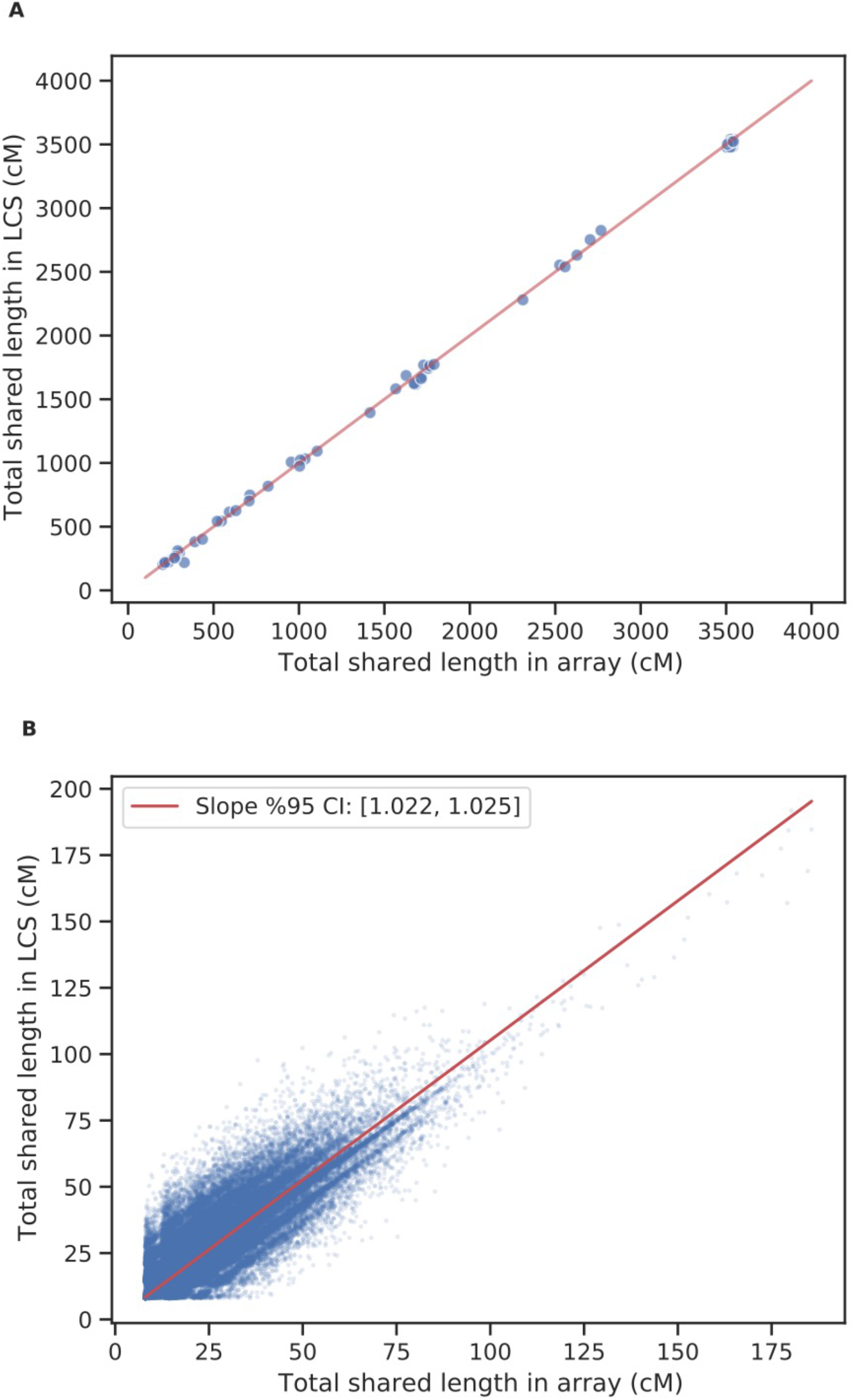
Regression of total matching pair length (cM) found for LCS vs array, for matches longer than 200 cM (A) and shorter than 200 cM (B).

### Detecting relatives with 1×LCS

Next, we measured the detection performance of finding relatives with 1×LCS compared to arrays as a function of the total IBD sharing. We started by measuring the ability of 1×LCS to recall the same relatives that were detected by the arrays. The missed relatives by 1×LCS are those who were assigned a total IBD=0 since the IBD detection pipeline did not find any perfectly matching segments longer than 6cM. However, we note that the array results are far from being a “truth set”. We reported above that when considering trios that were genotyped using homogenous platforms, 20% of the matches are detected only by children and not parents. These results suggest that IBD from array data is littered with spurious IBD segments that are the result of a miscall rather than a shared ancestor. Therefore, the recall rates indicate the repeatability between 1×LCS and arrays more, rather than the actual sensitivity of 1×LCS to detect true relatives.

Our results show that 1×LCS perfectly recalls the same set of close (>200cM) relatives as the array technology and 60% of all relatives. In all the 19 samples, no matches over a total IBD threshold of 200cM were missed by any one of the technologies. Moreover, out of 169 relatives that were detected by the array technology with a total shared cM of at least 100cM, only 25 were missed by the LCS, demonstrating a reproducibility of 85% (**Figure 3**). Interestingly, 23 out of the 25 array matches missed by the LCS technology belong to a single admixed individual of Ashkenazi Jewish and Central Asian ethnicity (sample #8 in **Table 1**). We suspect it is possible the mixed-ethnicity had adverse effects on the imputation and the matching results. We also considered the reciprocal case and measured the rate of relatives detected by 1×LCS and not the arrays as a function of their total IBD. This rate showed a similar picture with perfect concordance in close (>200cM) relatives, 94.5% in relatives above 100cM and 75% for all relatives detected by 1×LCS. Taken together, our results underscore the ability of 1×LCS to nearly perfectly recapitulate array results of third cousins and closer, which are the main focus of genetic genealogy.

**Figure 3.**
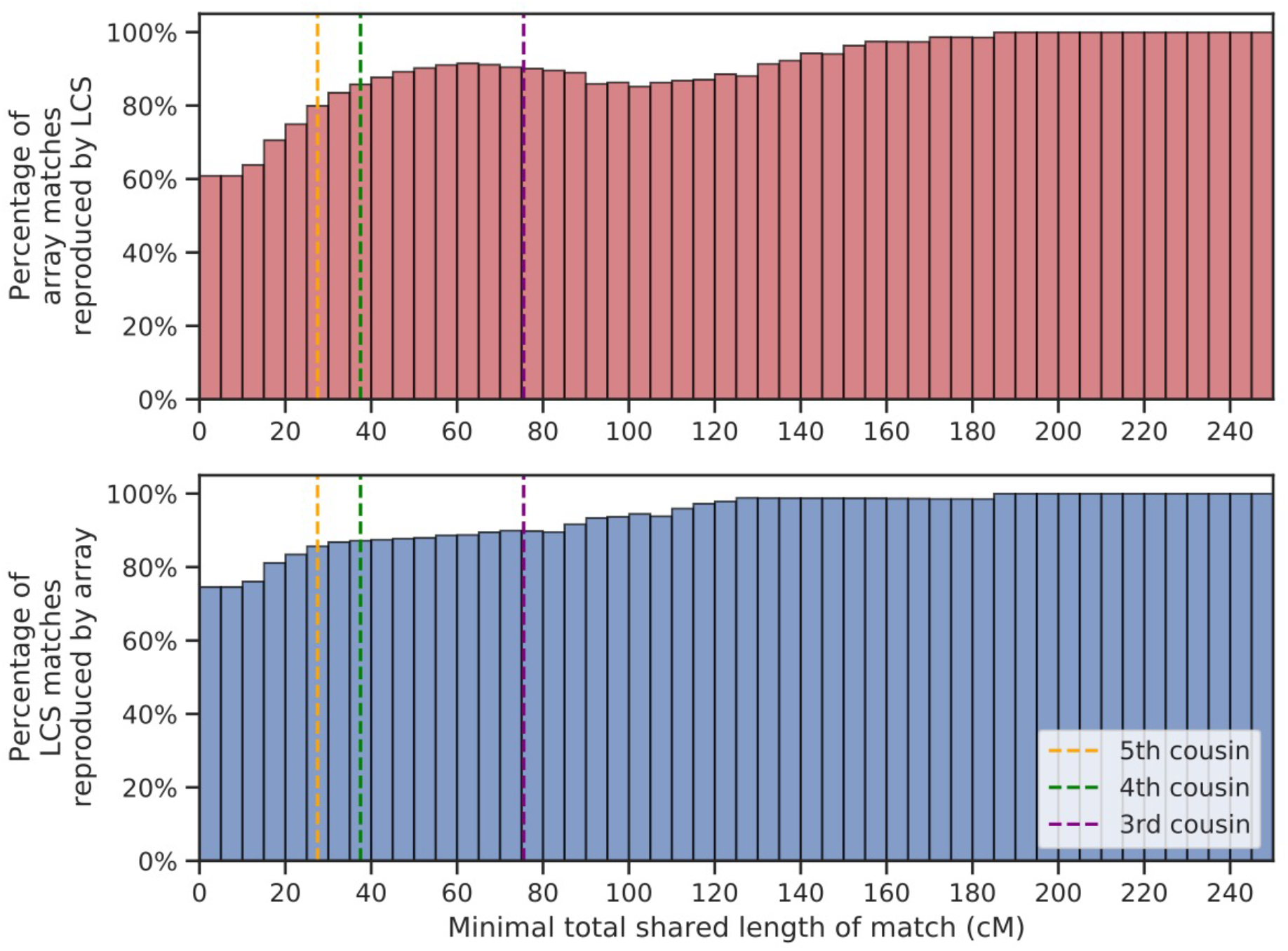
Percentage of array matches (red) and LCS matches (blue) replicated by the other technology. Each bin includes all matches with a total shared length of at least the bin's value. Above a relatedness level corresponding to 3rd cousins, matches are rarely missed by either technology. Vertical lines match the approximation of total shared IBD length (cM) for 5th, 4th and 3rd cousins (Bettinger & Perl 2018).

To better contextualize the performance of 1×LCS versus arrays, we focused on one individual in our cohort who was genotyped with both Omni and GSA arrays on top of the 1×LCS sequencing. Since this participant has two array genotyping technologies, it is possible to evaluate the performance of 1×LCS while using the differences between Omni and GSA as a baseline.

Our results show that the recall of 1×LCS in our pipeline is within the recall range of the two array technologies. The IBD pipeline detected 28,670, 18,890, and 25,469 relatives for the Omni, GSA, and 1×LCS, respectively (**Figure 4**). 1×LCS detected 18,051 relatives that were also detected by the Omni array, yielding a 64.5% recall rate for 1×LCS over Omni. This value is better than the GSA that detected 15,577 relatives that were also detected by Omni, and yielded a 54.3% recall rate of GSA over Omni. In the reciprocal analysis, the detection rate of Omni over GSA was 73.8%. We also examined the rate of relatives that are unique for each technology, which could be used to estimate the number of spurious relatives. The rate of unique relatives for 1×LCS was a bit higher than the arrays, with 23.0% compared to 21.2% for Omni and 11.57% for GSA. Again, these results show that the performance of 1×LCS to detect relatives using our pipeline is quite similar to the performance of finding relatives using heterogenous array technologies.

**Figure 4.**
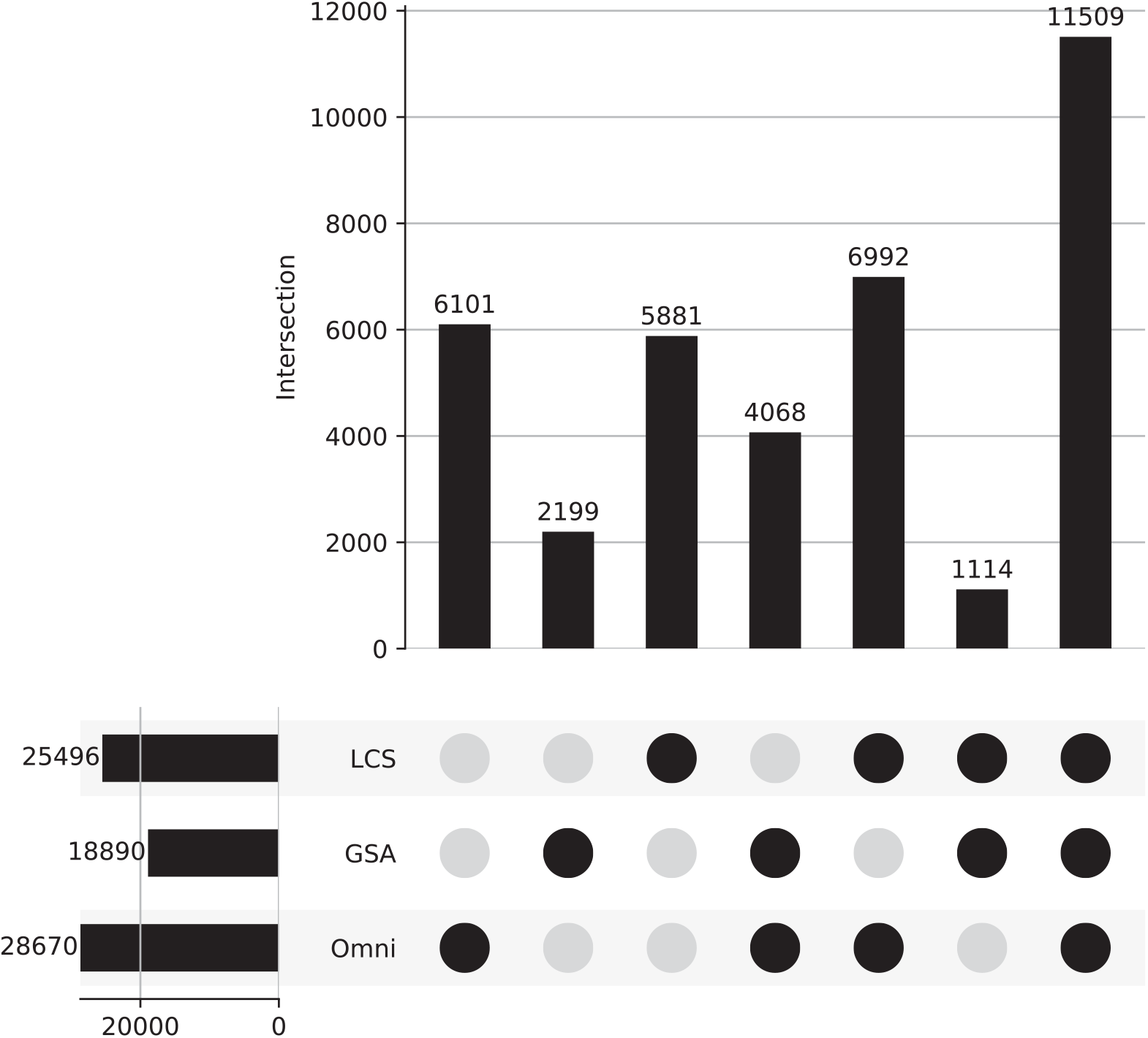
All matches found for sample #8, mapped to the groups of technologies by which they were detected. Most of the matches were detected by all three technologies.

## Discussion

Low coverage sequencing has received increasing attention in the last few years for various applications. In this work, we described the applicability of 1×LCS for relative matching, one of the main features of consumer genomics and the cornerstone for investigative genetic genealogy. Our results show that even when using an improved imputation panel, 1×LCS has a higher rate of genotyping discordance compared to datasets consist of homogenous array platforms but similar to datasets consist of heterogenous array platforms. To better tune the IBD pipeline for this higher discordance, we developed a hybrid IBD detection strategy and scanned a large set of known relatives that were genotyped using heterogenous array platforms. Using these relatives, we optimized the error tolerance of our hybrid pipeline and showed that it only modestly increases the rate of spurious IBD segments. We then used the improved pipeline to find relatives between 19 1×LCS samples and a database of 3 million individuals. We found that our results can be comparable to those of traditional genotyping arrays. For matches above 200cM, which are of the main interest of genealogical investigations, the total length of IBD segments between relatives did not display any bias and was virtually indistinguishable. For more distant relatives, 1×LCS performed well and any discrepancies were not substantially different from the discrepancies between Omni and GSA, the two most popular array technologies in consumer genomics.

We envision that in the short term 1× coverage can be a good sweet spot for consumer genomics, balancing quality and price. Indeed, the demand side of the consumer genomics tests is highly sensitive to price offering. Therefore, cutting the costs of goods (CoGS) by reducing the coverage can reduce the price and increase the uptake of these tests. However, we hypothesize that further cuts will not have a dramatic effect. First, sequencing only reflects a portion of the CoGS, which includes a sample collection device, boxing, shipment, accessioning, DNA extraction, library preparation, and bioinformatics. Assuming that sequencing is responsible for 30% of the COGS, cutting the coverage to say about 0.8× (20% less), will only affect 6% of the final CoGS. Second, our analysis showed that the presence of even a single sequence read dramatically improves the accuracy of the 1×LCS genotypes. A large number of the errors were in sites that were not supported by a single sequence read. This means that substantial cuts to the coverage are likely to result with poor genotyping accuracy.

In the mid/long term, we speculate that further improvements are possible. Our results showed that the size of the imputation panel dramatically increased the accuracy of the 1×LCS genotypes. Generally, a larger imputation panel will be available as deep coverage WGS data is accumulated. However, we also envision that as LCS datasets are accumulated in millions by consumer genomics companies, it might be possible to leverage the previous LCS to better impute a newly arrived LCS dataset. If successful, this can create a positive feedback loop of improved results with every LCS dataset. Eventually, such large imputation panels might enable to sample only very small parts of the genome to accurately determine each haplotype. However, we also note that if sequencing costs will continue to decrease, the economic motivation for lowering the coverage will be smaller. The CoGS contribution of sequencing to the overall CoGS will be diminished and thus further optimizations will not dramatically move the needle.

One open question is the right moment for consumer genomics to move from array technologies to low coverage sequencing. This manuscript reported that IBD detection can be achieved with 1×LCS and previous work showed that ethnicity estimation (Pasaniuc et al. 2012; Li et al. 2020; Homburger et al. 2019; Rustagi et al. 2017) and PRS scores can be calculated with 1×LCS at the same quality of arrays. As such, besides reports based on ultra-rare variants, these lines of study show that 1×LCS covers all main applications in consumer genomics. Getting the actual price economics of 1×LCS and arrays is complicated since commercial deals are usually not publicly available and they typically depend on volume commitment. Nonetheless, we estimate 1×LCS to be around $30 based on the publicly available NHGRI reports for 1Mb sequencing costs at Baylor and the Broad Institute with Illumina NovaSeq S4. The main question is whether the unit economic of arrays are substantially higher to justify the transition, when also including library preparation costs, amortization of capital equipment, and bioinformatics costs for each technology.

However, LCS might offer other advantages beyond unit economics that should be taken into consideration. First, in the past decade, the field has witnessed a minimal amount of innovation in array technology. The explosion of innovation in high throughput sequencing can usher in new types of reports and assays to consumer genomics that are not available with arrays. Second, the array production market for consumer genomics is dominated by a single player. On the other hand, high throughput sequencing attracts a larger number of manufacturers and therefore can facilitate competition and a more efficient market. These advantages can move forward the consumer genomics field and provide better experience to the tens of millions of people who participate in it.

## Methods

### Sample collection

We collected samples using the regular MyHeritage test kits, which are based on HydraFlock sterile absorbent tipped applicator (25-3206-H). After collection of buccal cells, the tip of the applicator was immediately placed in the lysis buffer, which is part of the MyHeritage test kit. Each participant contributed two samples using this procedure in two time points, one that was used for the experiments with our array technology and one for LCS. The 19 participants are MyHeritage employees and are mainly of European, Central Asian and Ashkenazi ethnicity. All agreed to the MyHeritage terms and conditions that permit the analysis in this manuscript. The samples were shipped to the relevant labs in room temperature.

### Array experiments

The samples were genotyped using standard genome-wide genotypes arrys; 12 of the samples were genotyped with the MyHeritage Illumina OmniExpress array; 6 with the MyHeritage Illumina GSA array, and one with the 23andMe version 3 Illumina OmniExpress array.

The MyHeritage arrays were genotyped in the GeneByGene lab and the processed genotype files were analyzed. The 23andMe sample was submitted to the MyHeritage pipeline through a user upload of his downloaded array information. The genotype files were phased and imputed using the public Haplotype Reference Consortium (S et al. 2016) database using Eagle 2.0 (Loh et al. 2016) and Minimac3 (Das et al. 2016).

### Sequencing experiments

The samples were sequenced using DNBseq technology of Complete Genomes in the Hong Kong facility of BGI. The samples were sequenced with a 150bp paired-end kit to a depth of 3× (17 samples) 10× (1 sample) and 30× (1 samples). Then, we downsampled the FASTQ files to generate exactly 10 million pair-end 150bp reads, which amount to 3Gbases of sequencing data.

### Sequencing analysis pipeline

We submitted the pair-end FASTQ sequence files to the Gencove analysis pipeline. This automatically aligns the reads and calls the variants. The output is an imputed and phased VCF file, as well as a BAM file of the alignment. We experimented with both Gencove original imputation based on the 1000Genomes projects imputation panel, and the HRC panel that yielded enhanced performance. The VCF and BAM files were downloaded to MyHeritage using the Gencove CLI and continued processing using our relative matching pipeline.

### 1×LCS coverage assessments

Using the BAM file that was produced by Gencove for each sample, we calculated coverage and depth statistics for each sample using Samtools 1.10 (Li et al. 2009). We used the Samtools built in function ‘stats’ to calculate the number of reads mapped, and the number of base-pairs covered by different number of reads. The average autosomal coverage was obtained by dividing the sum of all autosomal sites depth calculated with Samtools “depth”, by the total length of the autosome calculated from the Samtools “view” header. For the VCF files outputted by Gencove, we used the metadata fields of RC and AC (representing the number of reference and alternative calls mapped to the site) to divide all sites into those with an imputed VCF result that is based on at least one read or is imputed based on no reads at all. For high quality imputation SNPs analysis, we relayed on the information field of GP, containing the posterior probability assigned to each of the three possible genotypes at the site (homozygous-reference, heterozygous and homozygous-alternative). In this analysis we removed SNPs where the maximum value of the probabilities in the GP field was lower than 0.9.

### Calculating concordance of genotype results

Our concordance calculations were performed on different subsets of sites as described in the Results section. Usually, both NRC and overall concordance were calculated. For these calculations we remove any sites where the reported genotype in the array does not match the reference or alt alleles of the VCF, and any sites that are not called in the array results. Since VCF files might have multiple records for the same chromosome and base-pair position with different alternate alleles, in case both of them have a match with the array genotype (happens only for homozygous reference array genotypes), only one record will be kept to avoid multiple counts of the same position and overweighting multi-allelic sites. The concordance value is the percentage of sites where the genotype is identical out of all the compared sites. The NRC value is based on also filtering out any sites where none of the technologies reported the alternative allele before executing the concordance calculation described above.

The analysis of concordance stratified by minor allele frequencies is based on values obtained from the gnomAD dataset (Karczewski et al. 2020).

### Relative matching pipeline

Our relative matching pipeline is an hadoop based implementation of the seed-and-extend algorithm of GERMLINE (Gusev et al. 2009). In our genotyping error permissive version described here, we output segments of at least 6cM allowing for 4 heterozygous errors and up to 4 homozygous errors. Later, we filter out any pairs that do not have at least a single error free segment of 6cM or more. Our algorithm later performs stitching of all matching segments of a pair – merging of any two consecutive segments that are only separated by a small segment of less than 6Mbps, that is likely to be introduced due to genotyping errors. Following this process, we calculate the pair’s total shared cM length and number of shared segments. If any matching pair share less than a total of 8cM, it is discarded.

### IBD Sharing Analysis

For the regression analysis, we considered all pairs of matched individuals across all samples. Each pair was assigned an independent and dependent variable values based on the pair’s total shared length (in cM) assigned by the array technology and the 1×LCS technology, respectively. We then calculated the linear regression parameters for the entire data set while forcing the intercept to 0. We next performed a 10K bootstrap analysis by resampling residuals to obtain confidence intervals for the slope.

## Acknowledgments

We thank Jieming Chu, Weibin Liu, Xin Jin, Siyang Liu, Min Jian, Haorong Lu and the rest of the BGI team for rapid processing of the sequencing datasets in their facility. We also thank MyHeritage participants for enabling scientific discoveries.

