## Supplemental Figures for "Relative matching using low coverage sequencing"

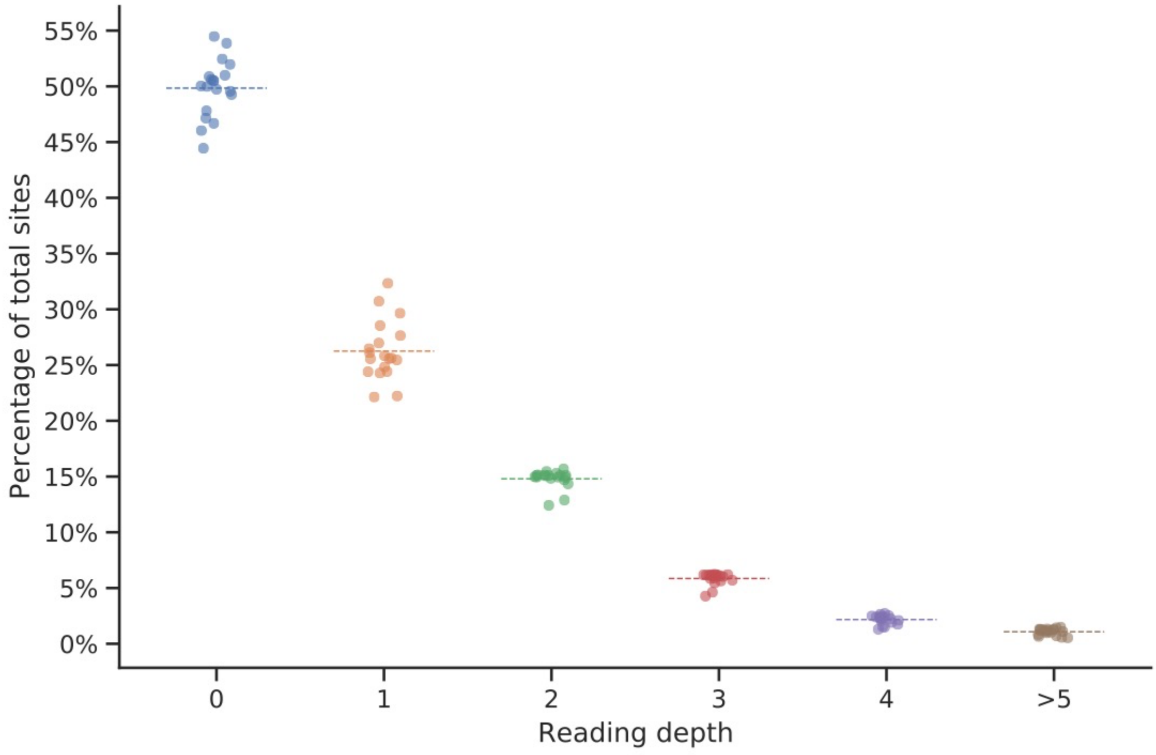


**Supplemental Figure 1:** Reading depth across sites of the entire genome (total 3,101,804,739 sites).

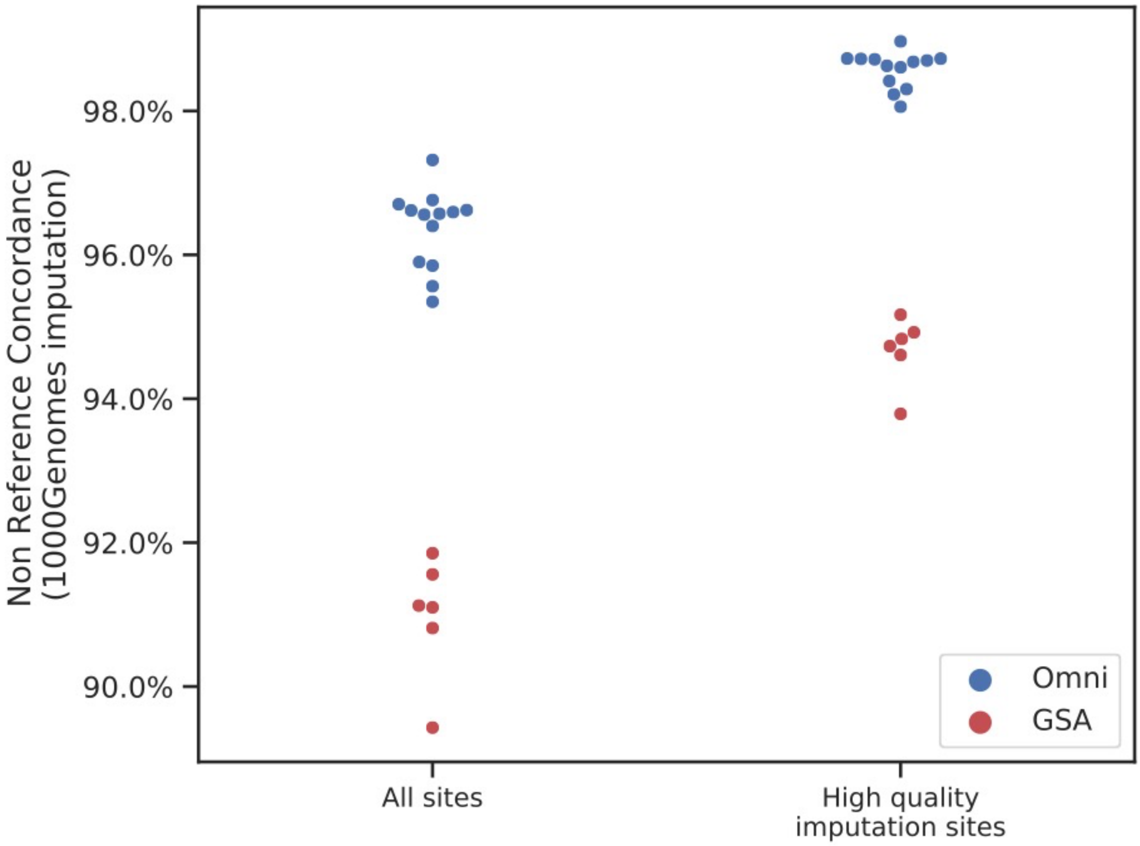


**Supplemental Figure 2:** NRC obtained for the 19 1xLCS compared to their array result, following imputation based on the 1000Genomes reference panel. Average NRC is 94.66%.
